# Visualizing *Borrelia burgdorferi* infection using a small molecule imaging probe

**DOI:** 10.1101/2020.09.04.284067

**Authors:** Madeline G Sell, David A. Alcorta, Andrew E. Padilla, Dakota W. Nollner, Nicole R. Hasenkampf, Havard S. Lambert, Monica E. Embers, Neil L. Spector

## Abstract

*In vivo* diagnostic imaging of bacterial infections is currently reliant on targeting their metabolic pathways, an ineffective method to identify microbial species with low metabolic activity. Here we establish HS-198 as a small molecule-fluorescent conjugate that selectively targets the highly conserved bacterial protein, HtpG (High temperature protein G), within *B. burgdorferi*, the bacteria responsible for Lyme Disease. We describe the use of HS-198 to target morphologic forms of *B. burgdorferi* in both the logarithmic growth phase and the metabolically dormant stationary phase. Furthermore, in a murine infection model, systemically injected HS-198 identified *B. burgdorferi* as revealed by imaging in post necropsy tissue sections. These findings demonstrate how small molecule probes directed at conserved bacterial protein targets can function to identify the microbe using non-invasive imaging and potentially as scaffolds to deliver antimicrobial agents to the pathogen.

---

Direct diagnosis of bacterial infections has historically relied on culturing microbes from blood or other body fluids and tissues. However, isolation of fastidious bacterial microbes, especially those that rapidly disseminate from circulation into tissues, including *Borrelia, Bartonella*, and *Leptospira* species, represent a diagnostic challenge [1]. Serologic immunoassays to detect host immune responses to stealth organisms that have mechanisms to evade the host response, such as *Borrelia burgdorferi*, the spirochete responsible for Lyme disease, lack sensitivity and accuracy especially in the setting of late stages of infection [2]. With an estimated prevalence of 340,000 new cases diagnosed every year in the United States, Lyme Borreliosis is the most common vector-borne illness in the U.S and Europe [3, 4].

A missed or delayed diagnosis of Lyme Borreliosis increases the risk of long-term morbidity related to chronic arthritis and neuropsychiatric symptoms [5]. Given the wide degree of variability in time of diagnosis and treatment, it has been reported that as many as 36% of individuals with Lyme Borreliosis will experience persistent symptoms [6]. Evidence from studies in animal models of Lyme Borreliosis and in infected humans points to persistent, active infection with *B. burgdorferi* as a probable underlying cause of the chronic symptoms in some individuals [7-10]. Other possible mechanisms include an exaggerated pathologic inflammatory response triggered by the presence of residual *Borrelia* antigens in the absence of active infection as the cause of persistent symptoms [11, 12]. One strategy to overcome the challenge of diagnosing stealth organisms, or organisms like *B. burgdorferi*, which exit the circulation soon after initial infection, is the development of imaging probes to visualize infection *in vivo*. Efforts on this front have been hindered to date by the lack of a probe that can specifically target a *Borrelia* protein that is highly abundant in all clinically relevant morphological states.

High temperature protein G (HtpG), the prokaryotic homologue of the 90 kDa mammalian heat shock protein (Hsp90), is an attractive target for diagnostic and potential therapeutic interventions for Lyme Borreliosis. HtpG is a highly expressed molecular chaperone found in most bacterial species [13]. We hypothesized it could be useful in detecting the presence of *B. burgdorferi* which is known for antigenic variation, loss of plasmids, and change in outer surface protein expression which hinder the ability to identify diagnostic protein targets [14]. Encoded on the linear chromosome of *Borrelia* rather than a plasmid, HtpG is conserved across different pathogenic species and strains of *B. burgdorferi* (Supplemental Fig 1) and is expressed throughout the different growth phases of *Borrelia*, which is essential when developing a targeted imaging probe [15]. Although less is known about the function of *Borrelia* HtpG, the eukaryotic homologue, Hsp90, is involved in maintaining intracellular proteostasis by regulating protein folding, trafficking, and preventing aggregation of denatured proteins [13, 16]. Amino acid sequence alignment from different prokaryotes and eukaryotic Hsp90, shows high conservation around the ATP binding regions (Supplementary Fig 2). However, the intervening amino acids between the conserved residues are often quite different, even across closely related prokaryotic forms. Exploiting these key inter-species differences between HtpG homologues could lead to an imaging diagnostic specific for *B. burgdorferi* and serve as a model for selectively imaging other pathogens. Imaging *B. burgdorferi in vivo* intermittently through the duration of treatment may also provide an objective and quantitative indicator of clinical response, and therefore guide the duration of therapy. We present a proof-of-concept study to demonstrate the feasibility of developing species selective small molecules against highly expressed HtpG for the diagnosis or treatment of stealth organisms by using the fluorescent small molecule, HS-198, as a prototype ligand.

## Results

### HtpG homologues exhibit varying drug binding affinities that can be exploited for selective drug targeting

Multiple sequence alignment of the N-terminal ATP domains of *B. burgdorferi* HtpG, *E. coli* HtpG, *Treponema denticola* HtpG, and Human Hsp90 shows significant overall homology, particularly at regions that make direct contact with ATP (Supplementary Figure 2) [17]. Subtle single amino acid differences are present within the catalytic clefts of these proteins, especially between residues that contact the nucleotide. Previous work in pathogenic fungus demonstrates that these differences may be exploited to create species selective agents [18]. We therefore reasoned that these differences may be sufficient to design small molecules that can similarly discriminate HtpG of stealth organisms. To test this hypothesis, we utilized a Fluorescent Linked Enzyme Chemoproteomic Strategy (FLECS) assay to test known Hsp90 inhibitors [19] for a binding to various HtpG constructs. First, we cloned and expressed in *E. coli* recombinant N-terminal GFP fusion forms of HtpG from *B. burgdorferi, E.coli* and *Treponema denicola*, the oral spirochete, as well as human Hsp90. We then incubated lysate after protein expression with an ATP sepharose resin to enable binding. In this assay, ATP is tethered to sepharose beads via its *γ*-phosphate enabling the nucleotide to bind to ATP-binding proteins such as Hsp90 and HtpG, but is non-hydrolyzable [20]. After proteins are bound to the ATP sepharose resin, small molecules with affinity for the ATP-binding site will cause elution, which can readily be detected and quantified via the GFP reporter. Identity of the eluted proteins were confirmed through an SDS-PAGE gel and further using MALDI mass spectrometry.

We measured protein elution as a function of drug concentration for PU-H71, Ganetespib, Radicicol, Geldanamycin, HS-198, HS-131, and HS-10 (Supplementary Fig 3 A, B, C, and D). These compounds were selected for testing based on their known binding to Human Hsp90 [21, 22]. To calculate the apparent dissociation constant (*K*_d app_) for each compound, we first calculated the Michaelis Constant (*K*_*m*_) of *B. burgdorferi, E. coli, and T. denticola* HtpG for ATP (supplementary Fig 1E) relative to the published human value (*K*_*m*_ = 300 *μM*), 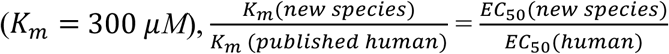, to be 181 μM, 226 μM, and 509 μM respectively [23]. Previously published results for *E. coli (K*_*m*_ of 250±82 *μ*M) are consistent with this method of analysis [24]. 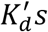 of all drugs tested in Supplementary Fig 3 were calculated using *K*_*d*_ = *EC*_50_/1+[Resin ligand/*K*_*m*_] and shown in Supplementary table 1. Several known Human Hsp90 inhibitors also bound to the three bacterial HtpGs demonstrating they lack species selectivity.

The binding of fluor-tethered Hsp90 inhibitor (HS-131), the non-tethered ligand analog (HS-10), and the inactive fluor-tethered analog N,N-dimethylamide (HS-198) to the recombinant proteins was also investigated (Figure 1A). In prior studies, we found HS-131 could localize to Hsp90 allowing for selective discrimination of human tumor cells exhibiting a malignant phenotype from non-transformed human epithelial cells [22]. HS-198 was designed as an inactive analog to HS-131 as the addition of methyl groups in place of hydrogens on the ligand prevented the molecule from binding to mammalian Hsp90 [22]. When the FLECS assay was performed with *B. burgdorferi* GFP-HtpG, both HS-131 and HS-198 selectively eluted the bound fusion protein from the ATP sepharose beads (Fig 1B). As expected from prior work, HS-131 effectively released recombinant human GFP-Hsp90*α* from the ATP sepharose beads while HS-198 did not (Fig 1C). HS-198 was the only inhibitor to target bacterial HtpG over Hsp90 (Figure 1 and Table 1). The method for calculating the *K*_d_’s were confirmed by comparison to the published results of Ganetespib [25]. HS-198 was unable to release *T. denticola* HtpG from the immobilized ATP resin, similar to human Hsp90. *E.coli* HtpG was released by HS-198 but exhibited minimally lower affinity than *B. burgdorferi* (Fig 1D). The selectivity of HS-131 was also found to show considerable variation across species (Fig 1E). In particular this molecule was completely inactive against *T. denticola* HtpG (Fig 1D). The finding that *T. denticola* is not recognized by HS-131 or HS-198 suggests that the latter probe can be used to diagnose the presence of *B. burgdorferi* over the related oral spirochete while not interacting with mammalian Hsp90.

**Table 1:**
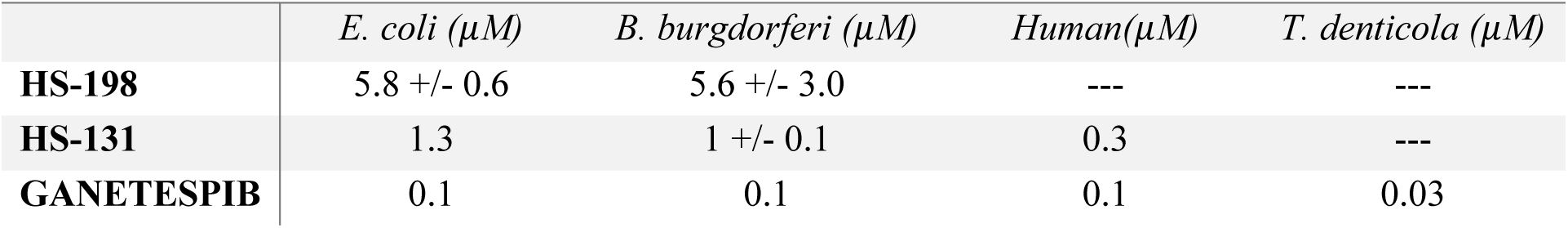
Kd’s of HtpG inhibitors. *K*_*d*_ = *EC*_50_/1+[Resin ligand/*K*_*m*_] (n=3) was utilized to calculate these values. The published value for Ganetespib (*K*_*d*_ = 110 *nM*) was used to confirm these calculations. Data shown as mean +/− SEM.

**Figure 1:**
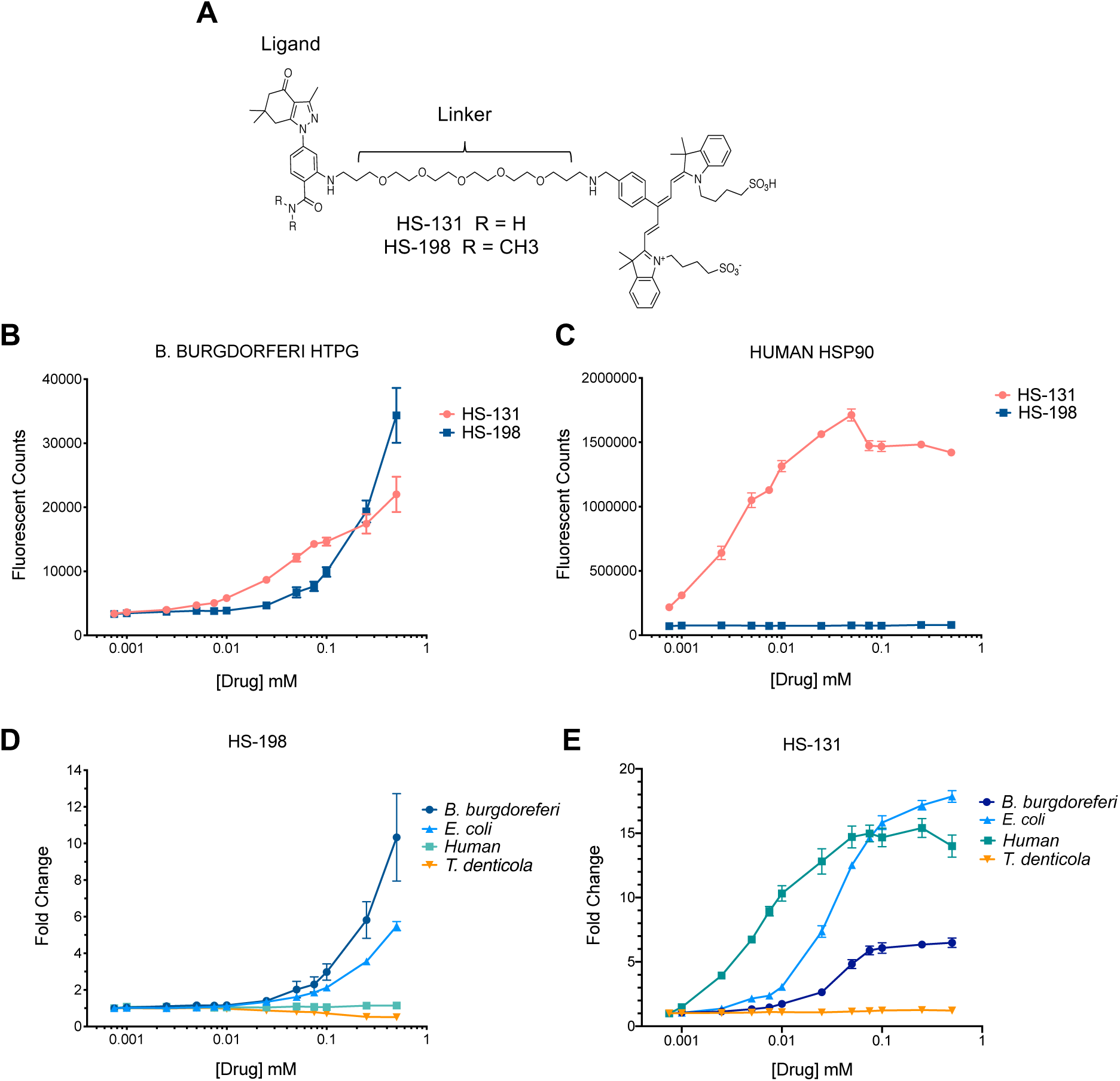
Structures and elution profiles of GFP-bound Hsp90 homologues with Hsp90 inhibitors (n=3). (A) Structures of HtpG inhibitors HS-198 and HS-131. (B) Fluorescent counts of drug induced GFP-bound *B. burgdorferi* HtpG elution. (C) Fluorescent counts of drug induced GFP-bound Human Hsp90 elution. (D) Fold change in HS-198 eluted fluorescent counts in *B. burgdorferi* HtpG, *E. coli* HtpG, human Hsp90 and *T. denticola* HtpG normalized to baseline. (E) Fold change in HS-131 eluted fluorescent counts in *B. burgdorferi* HtpG, *E. coli* HtpG, human Hsp90 and *T. denticola* HtpG normalized to baseline.

### HS-198 accumulates in *B. burgdorferi* in culture

We next investigated the selectivity of the fluorescent probe against live cultures of *B. burgdorferi*. HS-198 at 10 μM accumulates in *B. burgdorferi* in all of its morphological states including spirochetal, blebs, and aggregates (Fig 2A) as evidence by colocalization with DAPI. HS-198 was found to discretely bind the extra polymeric substance exported by *B. burgdorferi* as demonstrated by co-staining with Wheat Germ Agglutinin (WGA), a lectin that binds to N-acetyl-D-glucosamine, commonly used to identify extracellular matrices (Fig 2A). The video (Supplemental figure 4) of live *B. burgdorferi*, stained with HS-198 and visualized by CY5 (HS-198 signal) demonstrates that incubation with HS-198 overnight does not appear to affect *B. burgdorferi* morphology or motility, suggesting that the viability of the bacteria was not affected by the binding of HtpG. When incubated for 1 hour with 10 µM of HS-198, we routinely observed HS-198 uptake in >90% of *B. burgdorferi* spirochetes. To determine the nature of the zones of HS-198 accumulation, *B. burgdorferi* spirochetes were co-stained with primary flagellin antibody (Fig 2B). The flagellin antibody (red) rotates around HS-198 staining (blue). Since HS-198 must localize to an area within the flagellum rotation, this suggests that HS-198 accumulated inside the protoplasmic cylinder. Correlative Light Electron Microscopy (CLEM), a combination of fluorescent microscopy and high-resolution electron microscopy, confirmed separation from the outer membrane, and discrete spotting consistent with flagellum rotations blocking the signal, indicating localization of HS-198 inside the protoplasmic cylinder (Fig 2C).

**Figure 2:**
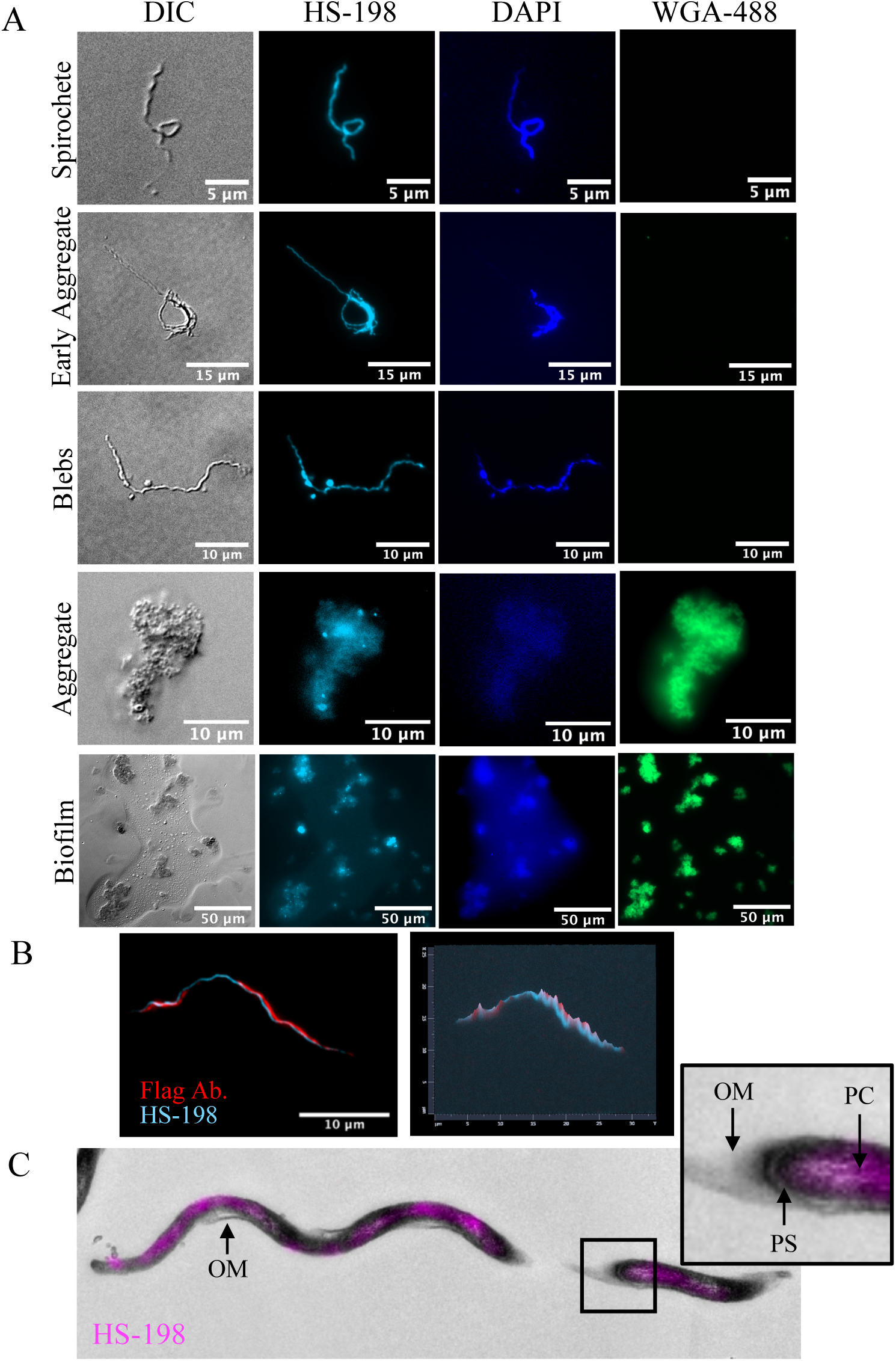
HS-198 accumulation and localization in *B. burgdorferi*. (A) Fluorescent imaging of *B. burgdorferi* in spirochetes, blebs, and aggregates stained with HS-198 (HtpG), WGA-488 (Extracellular Matrix), and DAPI (DNA) Images shown are representative of n=48. (B) Confocal images (100x) of HS-198 and Flagellin ab. staining of a spirochete n=5. (C) Correlative Light Electron Microscopy (CLEM) of HS-198 fluorescence overlaying an EM image of a *B. burgdorferi* spirochete. Outer Membrane (OM), Protoplasmic Cylinder (PC) and Periplasmic Space (PS) are identified. Images shown are representative of n=4 spirochetes with TEM sections of 65 nm.

### Imaging HS-198 in a mouse model of *B. burgdorferi* infection

To test the feasibility of using HS-198 to detect *B. burgdorferi in vivo*, we infected mice with approximately 500,000 spirochetes, and after 3 weeks, HS-198 (25 nmol) was injected via the tail vein. Six hours post injection, the animals were euthanized and tissue sections from the ear, tibiotarsal joint, and spleen, areas where *B. burgdorferi* is known to localize in this model, were prepared [9]. The sections were also co-stained with WGA and an anti-*B. burgdorferi* antibody. Examination of the sections by fluorescence microscopy identified *B. burgdorferi* in early and late stage aggregates in the ear and tibiotarsal joint (Fig 3A and B). Individual spirochetes were identified through HS-198 fluorescence as well as with anti-*B. burgdorferi* antibody (Fig 3C). Additionally, infected ear tissue from a mouse injected with HS-198 was co-stained with goat anti-rabbit conjugated with FITC as a non-specific antibody control to demonstrate the specificity of HS-198 *in vivo* (Fig 3D). These results demonstrate HS-198 selectivity for *B. burgdorferi* HtpG and localization to sites of *B. burgdorferi* infection in mice.

**Figure 3:**
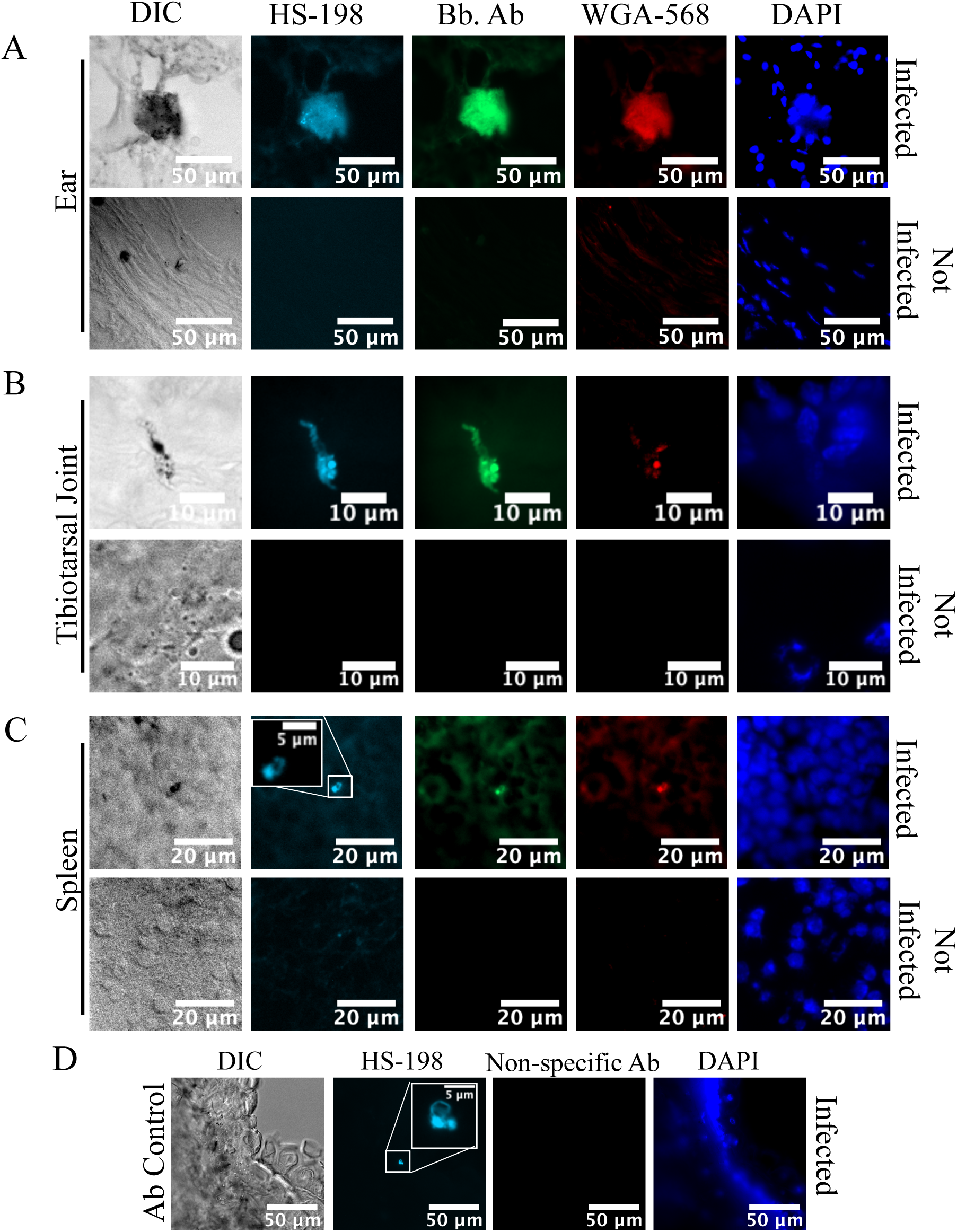
Fluorescence of tissues from mice infected with *B. burgdorferi* for 3 weeks and injected with HS-198 6 hours before sacrifice. (A) Ears (B) Tibiotarsal Joint and (C) Spleen were stained with FITC anti-*B. burgdorferi* antibody, WGA-568 (Extracellular Matrix), and DAPI (DNA). (D) Ears from mice injected with HS-198 were subsequently stained with goat anti-rabbit conjugated with FITC as a non-specific antibody consistent with the anti-borrelia antibody species. Images shown are representative of n=40 infected tissues and 15 uninfected.

## Discussion

Here we described an imaging agent able to exploit differences in the ATP binding site of a highly conserved, and highly expressed, heat shock protein, HtpG, to selectively target the stealth organism *B. burgdorferi* for a rapid diagnostic test of infection. HS-198 was previously developed as an inactive control molecule to demonstrate the selectivity of HS-131 to mammalian Hsp90 [22]. Fortuitously, when tested against purified recombinant *B. burgdorferi* HtpG, we discovered that the probe binds competitively to the proteins ATP binding site (Fig 1). The co-crystal structure of human Hsp90 with SNX2112 (an analog of the ligand portion of HS-131) shows an essential contact between D93 in the ATP binding site and the N2 nitrogen of the inhibitor [26]. The N,N-dimethylamide HS-198 analog disrupts the binding of the molecule to D93 of human Hsp90 [22]. We therefore attribute the elution of *B. burgdorferi* GFP-HtpG with HS-198 as being due to distinct amino acid differences in the ATP binding pocket that can accommodate the N,N-dimethylamide. Testing HS-198 against recombinant forms of *E. coli* and *T. denticola* HtpGs lends support to this hypothesis. In particular, this molecule was inactive against *T. denticola* HtpG, reflecting that even small substitutions within the ATP binding site of related prokaryotic HtpG molecules is sufficient to alter the specificity for these small molecules. These results also suggest that HS-198 can be used to diagnose the presence of *B. burgdorferi in vivo* over the related oral spirochete. However, *E. coli* demonstrated similar binding to HS-198. Studies in cattle have identified the colon as the site for *E. coli* persistence and proliferation [27]. As the colon is not a common location of *B. burgdorferi* localization, clinical presentation and infection localization would likely allow for differentiation between infections of *B. burgdorferi* and *E. coli*. However, future endeavors to modify the HS-198 ligand could improve species selectivity as well as increase the affinity.

We demonstrated that HS-198 is able to accumulate inside living *B. burgdorferi* at detectable amounts in all life stages, and once internalized, confocal and CLEM imaging showed HS-198 co-localized around the flagella, suggesting accumulation inside of the protoplasmic cylinder (Fig 2). This accumulation inside of the spirochete points to an opportunity to utilize the HS-198 ligand as drug lead for a payload delivery of toxins or antibiotics. Linezolid, for example, has maintained antimicrobial activity when linked to a fluorescent probe [28]. Linezolid was also found to be in the top 20 most effective drugs against *B. burgdorferi* in a high throughput screen of new drug candidates [29]. *B. burgdorferi* uptake of linezolid could possibly be improved by linkage to HS-198 ligand. Finally, imaging of tissues from mice that were inoculated with HS-198 while alive found that HS-198 is able to disseminate, localize to, and identify active *B. burgdorferi* infection (Figure 3).

There is an unmet need for selective imaging agents that could non-invasively detect infection with sparse organisms that are difficult to locate and culture. Current methods primarily include cell culturing, histological examination, and serological assays. Culturing methods prove problematic as the hallmark of fastidious organisms is that they are difficult to culture. In the case of *B. burgdorferi*, culturing only yields a 3.1% recovery rate from blood, or 45% recovery rate from erythema migrans rashes, which are only present in the initial stages of infection [1]. For detection using histological samples, invasive tissue biopsies are required, and there is a greater likelihood of scarce bacteria, such as *B. burgdorferi*, to be present but not in the location of the sample. Serological assays detect the presence of host antibodies to the bacteria. Cross-reactivity is a hurdle for diagnosis; for example patients with *B. quintana* infection may possess antibodies that are cross reactive with *Ch. Pneumonia, Chlamydia trachomatis*, and *Ch. psittac* [30]. Due to sequestration of antibody in antibody-antigen complexes and fluctuation in an antibody response, these tests become less inaccurate as the duration of infection increases which then often leads to false negative results, especially in the case of *B. burgdorferi* [31]. Targeting the host response for stealth organisms designed to evade the immune response is not an adequate means to assess the presence or absence of the organism. Methods to identify the etiology of the persistent symptoms in chronic Lyme Borreliosis are currently lacking as there are no non-invasive diagnostics available to determine whether there is an active infection or antibodies and remnants from a previous infection.

Direct detection of the pathogen confirms infection, and for this reason many researchers are turning to the development of imaging probes to visualize infection *in vivo*. These probes would allow whole body visualization and unlike select tissue samples they would provide the ability to monitor disease during and after treatment. To date, imaging strategies have largely targeted metabolic pathways utilized by bacteria, including the maltodextrin transporter expressed in gram-negative and positive-bacteria and 2-[18F]-fluorodeoxysorbitol that utilizes sorbitol, a metabolic substrate for *Enterobacteriaceae* [32, 33]. In addition, fluorine-18-fluorodeoxyglucose (18F-FDG) positron emission tomography (PET) has been used to identify foci of increased glucose uptake, including inflammation, infection and malignancies [34]. Although targeting metabolic pathways may be effective in actively replicating bacteria with high metabolic requirements, persistor and stationary growth phase bacteria exhibit low metabolic activity and are therefore not amenable to this approach [35]. Importantly, none of these imaging approaches are capable of Lyme diagnostics.

To solve this problem, we identified a protein that is expressed at high levels in all growth phases and morphological variants of *B. burgdorferi*, and, in order to identify persisting dormant bacteria, one that is not dependent upon active metabolism. Like its mammalian counterpart, Hsp90, HtpG is highly abundant at levels identifiable in the serum of infected patients, and by virtue of its ATP binding pocket, highly druggable. Although optical imaging probes tethered to HtpG have utility in histological and cellular studies, PET enabled versions of tethered selective small molecules may offer a non-invasive approach to detect unresolved *B. burgdorferi* infections by whole body imaging. Our discovery that HS-198 is a *B. burgdorferi* selective probe that labels the bacteria both in culture and mice has broad implications for the development of a Lyme diagnostics or treatment. Future developments of HS-198 include a) agent detection using in vitro imaging systems (IVIS) that would allow imaging of a whole live mouse to identify active infection, b) modifications to the ligand to increase specificity and affinity, c) an exchange of the fluorescent molecule for a PET agent for imaging, and d) combining the HS-198 ligand with a toxin or antimicrobial agent to selectively deliver payloads to the bacteria for eradication.

## Online Methods

### GFP-fusion Protein Plasmid Construction

The *B. burgdorferi, E.coli, and Treponema denticoli* HtpG DNA sequences were PCR cloned from genomic DNAs using primers designed to allow in-frame ligation independent cloning with GFP in the IPTG inducible bacterial expression vector, 1GFP (pET His6 GFP TEV LIC cloning vector (1GFP) was a gift from Scott Gradia (Addgene plasmid # 29663). Human Hsp90-alpha was similarly cloned from cDNA generated from the human breast epithelial line, Mcf10A. Fusion protein cloning was done using the ligation independent cloning procedure via single nucleotide T4 DNA polymerase treatment of vector and purified PCR products then annealing of complementary single strand ends before transformation into DH5 *E. coli*. Proper fusion junctions and fidelity of HSP inserts were confirmed by sequencing then the plasmid was transformed into *E. coli* BL21(DE3) for protein expression. Fusion protein synthesis was induced by incubation with 1 mM IPTG or autoinduction [36]. Production of GFP-HtpG/Hsp90 was confirmed by identification of GFP and HtpG peptides with MALDI using a ABSciex 5800 TOF/TOF mass spectrometer in the induced band identified on a SDS-PAGE gel. To complete binding curves, several liters of induced cultures were generated, pelleted with centrifugation, flash frozen in liquid nitrogen and then stored at −80 °C until subsequent FLECS analysis.

### FLECS Assay

γ-Linked ATP Sepharose matrix was generated as described previously by the Haystead lab [19]. BL21 bacterial pellets expressing GFP-fusion proteins of *E. coli, B. burgorferia, T. denticola* HtpGs and human Hsp90 were lysed with B-PER Complete Bacterial Protein Extraction Reagent (ThermoScientific 89821) and centrifuged at 35,000 rpm (Beckman coulter type 45 Ti Fixed Angle Rotor) to pellet remaining BL21 material. The clarified lysate was added to a column containing ATP bound sepharose. The column was washed with low salt buffer [150 mM NaCl, 25 mM of HEPES, pH 7.4, 1 mM of DTT, and 60 mM MgCl_2_], followed by high salt buffer [1M NaCl, 25 mM of HEPES, pH 7.4, 1 mM of DTT, 0.1% IGEPAL, and 60 mM MgCl_2_] and once again low salt buffer to remove unbound protein. Next, the resin with bound proteins (50 μl) was transferred to 0.2 μm polyvinylidene fluoride filter 96-well plate (Corning) sitting on top of a black flat-bottomed 96-well catch plate (Corning). Small molecules or ATP were added to each well (50 μl) and the plates were centrifuged using an Eppendorf Centrifuge 5810 at 1,100 rpm for 1 min. The eluted proteins were measured in a fluorescent plate reader to detect the GFP. As a means to confirm the presence of HtpG verses fluorescent compounds that could cause an assay signal, samples were analyzed by SDS-PAGE followed by silver stain and mass spectrometry.

### *B. burgdorferi* Bacterial Growth

*B. burgdorferi* strain B31 was obtained from ATCC (ATCC 35210). Cultures were grown in Barbour–Stoenner–Kelly medium (BSK-II) without gelatin [16] and supplemented with 6 % rabbit serum at 34 °C. Low passage cultures (less than 4) were utilized. Later stage morphological variants were obtained by increasing culture time to approximately 8 days to achieve a density of ∼10^8^.

### *B. burgdorferi* Bacterial Imaging

Bacterial cultures in logarithmic or stationary phase were incubated in media for 1 hour with 10 μM HS-198 and wheat germ agglutinin (WGA) conjugated with Alexa 488 at 10 µg per ml (Biotium Cat #29022), cells were then washed with fresh media and mounted with prolong Gold with DAPI. Samples were visualized using a Zeiss Axio Imager Upright microscope, ×62 oil objective, 405, 488 or 633 nm laser and DIC.

### CLEM

CLEM (Correlative Light Electron Microscopy) was performed by Shannon Modla and Jeff Caplan at the bioimaging center of the Delaware Biotechnology Institute. Cells were attached to an Ibidi µ-dish containing an imprinted 500 µm cell location grid using 4% PFA for 20 min. Cells were then incubated with 10 *μM* of HS-198 for 10 minutes followed by PBS. Cells were given to the Delaware institute for further processing and imaged using Zeiss 880 Airyscan. The alphanumeric pattern from the Ibidi µ-dish was imprinted on the freshly exposed surface of the resin, which allowed the same region of interest imaged by light microscopy to be re-identified in the ultramicrotome. Ultrathin serial sections were collected using a Leica UC7 ultramicrotome and picked up onto 2 x 1 copper slot grids, which were then dried on a domino rack (Electron Microscopy Sciences; Cat No 70621) coated with 0.5% formvar in ethylene dichloride. Serial sections were examined on a Libra 120 transmission electron microscope operating at 120kV, and images were acquired with a Gatan Ultrascan 1000 CCD using Gatan Digital Micrograph software. To capture TEM images of the entire bacterium, overlapping images were collected at 5000X and 16000X magnifications and then stitched together using the ImageJ plug-in MosaicJ.

### Immunohistochemistry

10 *μm* sections of frozen tissues were incubated with WGA (Biotium Cat #29077) for 1 hour in PBS. Tissues were washed and fixed with 3:1 methanol to acetic acid. Slides were incubated with FITC conjugated *B. burgdorferi* polyclonal antibody (Thermo Fisher Cat # PA1-73005) for 1hour at room temperature. Non-injected or infected tissues were washed and incubated with 1 μM of HS-198 for 30minutes as control, injected tissues were not incubated with HS-198. Slides were washed and mounted with DAPI prolong Gold. Imaged using a Zeiss Axio Imager Upright microscope, 62X oil objective, 405, 488 or 633nm laser and DIC. Images were analyzed using FIJI software.

### Animals, spirochetal inoculation, and treatment

Practices in the housing and care of mice conformed to the regulations and standards of the Public Health Service Policy on Humane Care and Use of Laboratory Animals, and the Guide for the Care and Use of Laboratory Animals. The Tulane National Primate Research Center (TNPRC) is fully accredited by the Association for the Assessment and Accreditation of Laboratory Animal Care-International. The Tulane University Institutional Animal Care and Use Committee approved all animal-related protocols, including the infection and sample collection from mice.

#### In vivo mouse model for tissue studies

Seven C3H/HeN (Charles River labs) mice were anesthetized with isoflurane gas, 1.5-2%. Mice were infected with 500,000 logarithmic stage *B. burgdorferi* strain N40 in < 0.5 ml sterile saline by subcutaneous injection in the nape of the neck via 25-gauge needle. Ear punch biopsies were collected from mice at 7 and14 days post-inoculation to confirm infection. Disposable 2mm punches were used on the outer rim of the ear to collect skin from anesthetized mice. The biopsies were placed in BSK-H (Sigma) culture medium and grown for 2 weeks to confirm infection. After 3 weeks, mice were injected with 25nmol/animal of HS-198 in the tail vein with a tuberculin needle. 6 hours later mice were euthanized by CO2 inhalation, followed by harvest and flash-freezing of tissues, including the ear skin, spleen and tibiotarsal joints.

## Acknowledgements

The authors would like to thank Dr. Ben Carlson and Dr. Yasheng Gao in Duke Microscopy core for assistance with confocal microscopy, Jennifer Miller at Galaxy for assisting in the design of our *B. burgdorferi* culturing method, and Shannon Modla and Jeff Caplan for their work with CLEM. We would like to thank Timothy Haystead and his lab members for discussions and advice on interpretation of FLECS assay data and mass spectrographic confirmation of the fusion protein identities as well as providing the HS-198 and HS-131 generated in the lab for these studies.

## Author Contributions

M.S and A.P performed the FLECS assays, M.S stained the *B. burgdorferi* and tissues, and performed all imaging. D.A, D.N. and A.P. cloned, expressed and validated all of the GFP-heat shock protein fusion proteins. M.E, N.H. and H.L. provided DNA, and provided the infected, HS-198 injected, tissues for additional processing. P.H. created and provided HS-198 and HS-131. MS, DA, and NS designed the experiments. MS, DA, and NS wrote the manuscript.

